# The subtle balance of insulin and thyroxine on survival and development of primordial follicles cultured *in vitro* enclosed in ovarian tissue

**DOI:** 10.1101/655589

**Authors:** Victor M. Paes, Laritza F. Lima, Anna-Clara A. Ferreira, Carlos H. Lobo, Benner G. Alves, Ana-Paula R. Rodrigues, Ariclecio C. Oliveira, Jose R. Figueiredo, Jean M. Feugang

**Author notes:** Corresponding author (JMF). Department of Animal and Dairy Sciences, Mississippi State University, Starkville, Mississippi, United State of America (Jean M. Feugang). These authors contributed equally to this work (Victor M. Paes, Jose R. Figueiredo, Jean M. Feugang). These authors also contributed equally to this work (Laritza F. Lima, Anna-Clara A. Ferreira, Benner G. Alves, Ana-Paula R. Rodrigues, Ariclecio C. Oliveira).

## Abstract

Thyroid hormones have presented a positive hormonal interaction on follicular development of secondary follicles and oocytes from antral follicles; however, the effect of thyroid hormones on primordial follicles is unclear. Here we investigated the *in vitro* effects of combined insulin and thyroxine on caprine primordial follicle survival and development. Ovarian tissues were cultured for 1 or 7 days using 10 ng/ml (low) or 10 µg/ml (high) insulin in the absence or presence of thyroxine at 0.5, 1 or 2 µg/ml. Thereafter, follicular survival and development, gene expression related to apoptosis (*Bcl2/Bax*), insulin and thyroid receptors, and estradiol and reactive oxygen species production were evaluated. In low-insulin conditions, supplementation with 2 µg/ml thyroxine maintained follicular survival similar to non-cultured control, while 0.5 µg/ml thyroxine enhanced the survival (P<0.05) in comparison to thyroxine-free treatment. Only treatments containing low-insulin and thyroxine at 0.5 or 2 µg/ml increased (P<0.05) reactive oxygen species production from day 1 to day 7. Contrarily to high-insulin containing medium, the presence of thyroxine in low-insulin medium yielded higher stromal cell density (P<0.05). There were higher (P<0.05) estradiol production and *Bcl2/Bax* ratio in low-insulin versus high-insulin treatments on day 1 and 7, respectively. High levels of both insulin and thyroxine showed better follicular development (P<0.05), yielding great follicle and oocyte diameter. Finally, the high-insulin level led to insulin and thyroid receptors expression reduction as compared to non-cultured control. In conclusion, the combination of low concentrations of insulin and thyroxine better maintained follicle survival, while high levels ensured better follicular development.

## Introduction

Mammalian ovarian folliculogenesis is a complex process regulated by autocrine, paracrine, and endocrine factors (1). Regarding endocrine signaling, both thyroid hormones (TH) triiodothyronine (T3) and thyroxine (T4) have a role-play on follicular development (2). In this sense, an imbalance of TH such as hypo- or hyper-thyroidism has been associated with infertility in woman (3). The majority of released TH is in the form of T4, which it is regulated by a subtle modulation of deiodinase enzymes (DIO) present in several tissues. DIO thus could inactive T4 (DIO I and III), or to convert T4 to T3 (DIO I and II) to exert its genomic actions through interaction with its nuclear receptor (4, 5). Due to its importance, thyroid receptor (TR) has been detected in several species (6). Although caprine species have been described as a good model to study human ovary (7-9), the presence of TR in the caprine ovary is unknown.

The disturbs of the TH in murine causes an imbalance in both redox homeostasis (10) and estradiol levels leading to follicular apoptosis (11). To better understand about the effects of TH at the cellular level, *in vitro* experiments have been performed either using T3 (12-14) or T4 (15-19). The use of T3 and/or T4 in association to other hormones enhance survival and development of secondary follicles in murine (12, 14), and ovine (17), as well as bovine embryo development (13, 16). Thyroxine concentrations usually used in the experiments with preantral follicles vary from 0.5 µg /ml to 2 µg/ml (15, 17-20). Nevertheless, even though few in vitro studies reported the effects of TH on the development of late preantral follicle categories to the best of our knowledge of its in vitro role on primordial follicle survival and development remains unknown.

Among the various hormones that act on folliculogenesis, insulin has an important role in follicle development due to its action on cell survival and proliferation (21). Our team has shown that insulin effects could be influenced by follicular category (22, 23), as well as its interactions with other substances (24-26). For instance, the use of insulin at 10 ng/ml during *in vitro* culture of either primordial (22), secondary or tertiary follicles (23) yielded greater oocyte growth than using 10 µg/ml insulin. On the other hand, secondary follicles cultured *in vitro* with insulin at 10 µg/ml in association to FSH (24) and GH (25), or FSH + VEGF (26) enhance meiosis resumption in goat oocytes. Although the hormonal network is a complex process, the balance of insulin and thyroxine concentration on in vitro primordial follicles development is unclear.

Therefore, the novelty of the present study was to investigate the *in vitro* effects of combined insulin and thyroxine on primordial follicles evaluating: i) follicular survival and development; ii) Estradiol and ROS production; iii) quantitative expression of some genes related to apoptosis (*Bcl2/Bax*), and hormone receptor for both thyroid (TR) and insulin (IR).

## Material and Methods

### Ethics statement

All procedures were approved by Ethics Committee in Animal Experimentation of State University of Ceará (2789214/2017).

### Source of ovaries

Ovaries (n=16) from 8 adult mixed-breed goats were collected from a local slaughterhouse. Immediately after slaughter, ovaries were removed and washed once in 70% alcohol and twice in minimum essential medium (MEM) with HEPES (MEM-H) and antibiotics (100 µg/ml penicillin and 100 µg/ml streptomycin). The ovaries were transported to laboratory within 1 hour in MEM-H at 4°C (27). All chemicals used in the present study were purchased from Sigma Chemical Co (St. Louis, MO, USA) unless otherwise indicated.

### Experimental design

In experiment I (n=10 ovaries), the effect of different insulin and thyroxine combinations on the *in vitro* survival and development of goat preantral follicles enclosed in ovarian tissue was investigated (Fig 1). The experiment II (n=6 ovaries) was conducted to evaluate the effects of insulin and thyroxine on preantral follicles at the molecular level.

**Fig 1.**
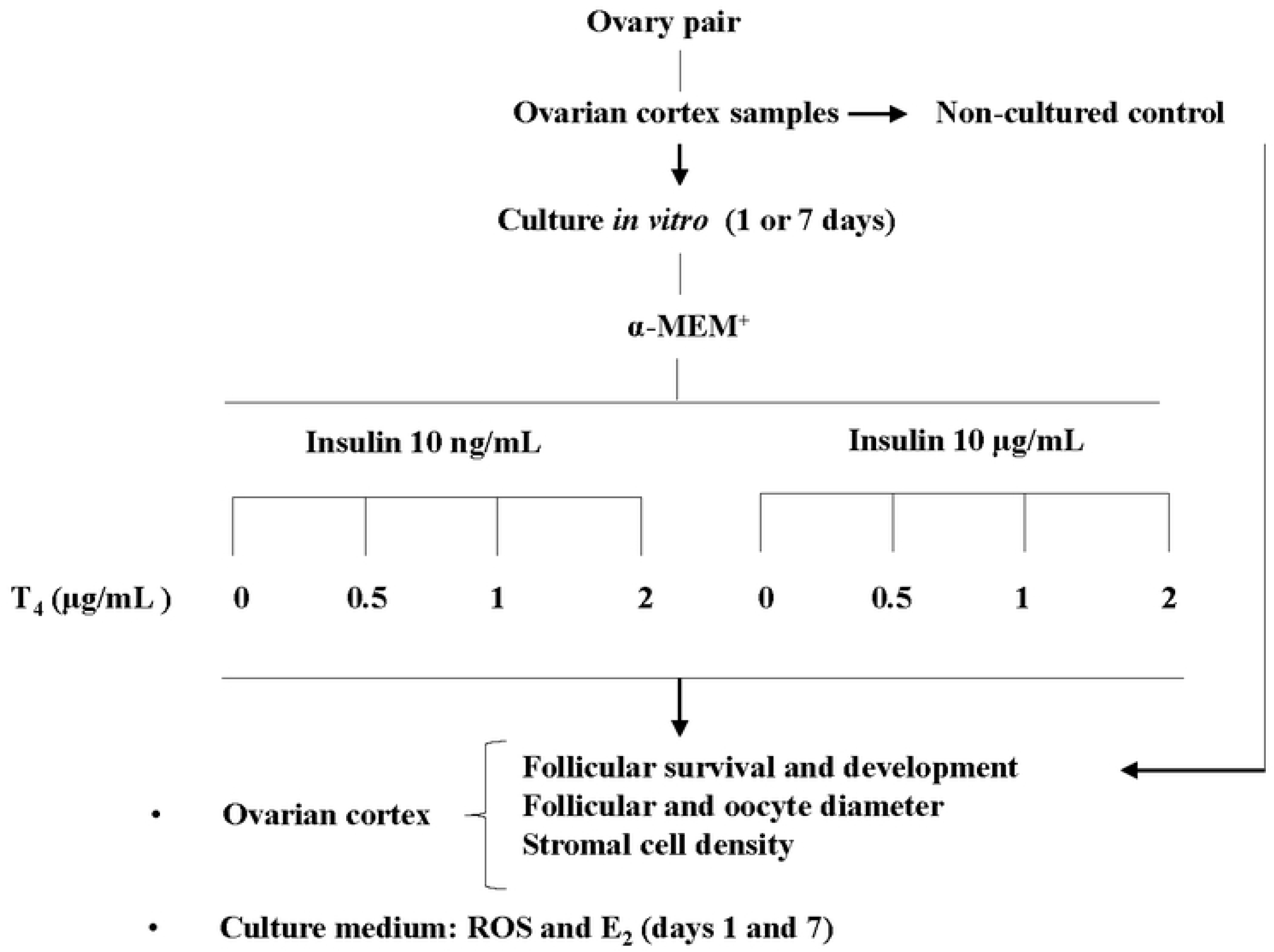
Schematic of the experimental design.

For both experiments, ovarian cortex samples from each ovarian pair were cut into slices approximately 3 × 3 × 1 mm in size using a needle and scalpel under sterile conditions. For each animal, one slice of tissue was randomly selected and immediately fixed (non-cultured control, day 0). The remaining slices of ovarian cortex were cultured individually in 1 ml of culture medium in 24-well culture dishes for 1 (Experiment I) or 7 day (Experiment I and II), in an atmosphere of 5% CO_2_ in air at 39°C. The basic culture medium consisted in α-MEM supplemented with 1.25 mg/ml BSA, 5.5 µg/ml transferrin, 5.0 ng/ml sodium selenite, 2 mM glutamine, 2 mM hypoxanthine, 100 µg/ml penicillin and 100 µg/ml streptomycin (22). The ovarian slices were distributed to the following treatments: Low or high concentration of insulin (10 ng/ml and 10 µg/ml, respectively) alone or associated with thyroxin at different concentrations (0.5, 1.0 and 2.0 µg/ml). Each treatment was repeated 5 times for Experiment I and 3 times for Experiment II, using ovaries of different animals. The culture medium was stabilized at 39°C overnight prior to use and was replenished every other day. Culture media obtained on day 1 and day 7 were stored at −20°C for subsequent assays.

## Experiment I

### Morphological analyses and assessment of in vitro follicular growth

Ovarian tissues from all treatments (non-cultured control or cultured for 1 or 7 days) were fixed in Carnoy’s solution for 12 h, dehydrated in increasing concentrations of ethanol, cleared in xylene, embedded in paraffin wax (Synth, São Paulo, Brazil) and serially sectioned at 7 µm. Every section was mounted on glass slides and stained with hematoxylin and eosin (HE). Follicle stage and survival were assessed microscopically on serial sections. Coded anonymized slides were examined by microscopy (Nikon, Sendai, Japan) at 400x magnification. For each treatment, at least 30 preantral follicles were analyzed per animal (replicate). Only preantral follicles with a visible oocyte nucleus were counted. The follicles were considered degenerated if they contained a pyknotic oocyte nucleus or shrunken ooplasm, with or without disorganized granulosa cells and/or detachment of the basement membrane (27).

According to their developmental stage, preantral follicles were defined as primordial (oocyte surrounded by one layer of flattened pregranulosa cells) or developing, with the latter category further divided into intermediate (oocyte surrounded by one layer of flattened pregranulosa and cuboidal granulosa cells), primary (oocyte surrounded by one layer of cuboidal granulosa cells) or secondary (oocyte surrounded by two or more layers of cuboidal granulosa cells) (27).

To evaluate follicular activation (transition from primordial to growing follicles, when surrounding squamous pregranulosa cells become cuboidal and begin to proliferate) and growth, only morphologically normal follicles with a visible oocyte nucleus (equatorial section) were recorded, and the proportion of primordial and growing follicles was calculated at day 0 (non-cultured control) and after 1 or 7 days of culture with treatment. In addition, follicular and oocyte diameters were measured using Nikon NIS-Elements software (Nikon NIS-Elements 3.0, 2008, Nikon, Tokyo, Japan), considering the major and minor axes of each follicle and oocyte. The average of these two measurements was used to determine the diameters of both the oocyte and the follicle (28).

### Ovarian stromal density

The density of ovarian stromal was evaluated by calculating the stromal cell per 100 µm^2^. For each treatment, ten fields per slide were assessed and the mean number of stromal cell per field was calculated (29).

### Hormone assays

To evaluate follicular steroidogenesis *in vitro*, concentrations of estradiol (E2) were measured in spun culture media at day 1 and day 7 to reflect a 24-h hormonal production. Standard dilutions were made according to manufacturer’s instructions using competitive immunoassay commercial kit (enzyme linked fluorescence assay VIDAS, Biomerieux, Marcy l’Etoile, France). The analytical sensitivity was 9 pg/ml, ranging from 9 to 3000 pg/ml, while the intra-assay coefficient of variation was 5%.

### Levels of reactive oxygen species (ROS)

Ten microliters of medium were incubated with D-glucose (1 mg/ml), SOD (100 U/ml), HRP (0.5U/ml) and Amplex Red (50 μM), as previously described (30). The fluorescence was measured in a microplate reader (Victor X4), using excitation at 530 nm and emission at 595 nm. The concentration of reactive oxygen species was expressed as micromoles of H_2_O_2_.

## Experiment II

### RNA extraction and real-time PCR (qPCR)

For RNA isolation, three ovarian fragments were collected from non-cultured control and each experimental group. The samples were collected in microcentrifuge tubes (1.5 ml) on ice and stored at −80°C until RNA extraction. Total RNA was isolated and purified with Trizol® reagent method (Invitrogen, Carlsbad, CA, USA) according to the recommendations of the manufacturer and further purified with PureLink ™ RNA Mini Kit (Ambion®, Carlsbad, CA, USA). The isolated RNA preparations were treated with DNase I and Pure LinkTMRNA Mini Kit (Invitrogen, São Paulo, SP, Brazil). Complementary DNA was synthetized from the isolated RNA using SuperscriptTM II RNase H-reverse Transcriptase (Invitrogen, São Paulo, SP, Brazil). The quantitative polymerase chain reaction (PCR) was carried out in a final volume of 20 µL containing 1 µL of each complementary DNA, one × Power SYBR Green PCR Master Mix (10 µL), 7.4 µL of ultra-pure water, and 0.5 µL (final concentration) of both sense and antisense primers. Primer sequences for target genes are shown in Table 1. The reference gene glyceraldehyde 3-phosphate dehydrogenase (GAPDH) was selected as an endogenous control for normalizing gene expression among the samples. Primer specificity and amplification efficiency were verified for each gene. The PCR conditions consisted of an initial step for denaturation and polymerase activation for 15 minutes at 94°C, followed by 40 cycles of 15 seconds at 94°C, 30 seconds at 60 °C, and 45 seconds at 72°C. A final extension was performed for 10 minutes at 72°C. The specificity for each primer set was tested with melting curves that were carried out between 60°C and 95°C for all genes. All amplifications were carried out in a Bio-Rad iQ5 (Hercules, CA). The 2^-ΔΔCT^ method was used to transform threshold cycle values into normalized, relative expression levels (31).

**Table 1.**
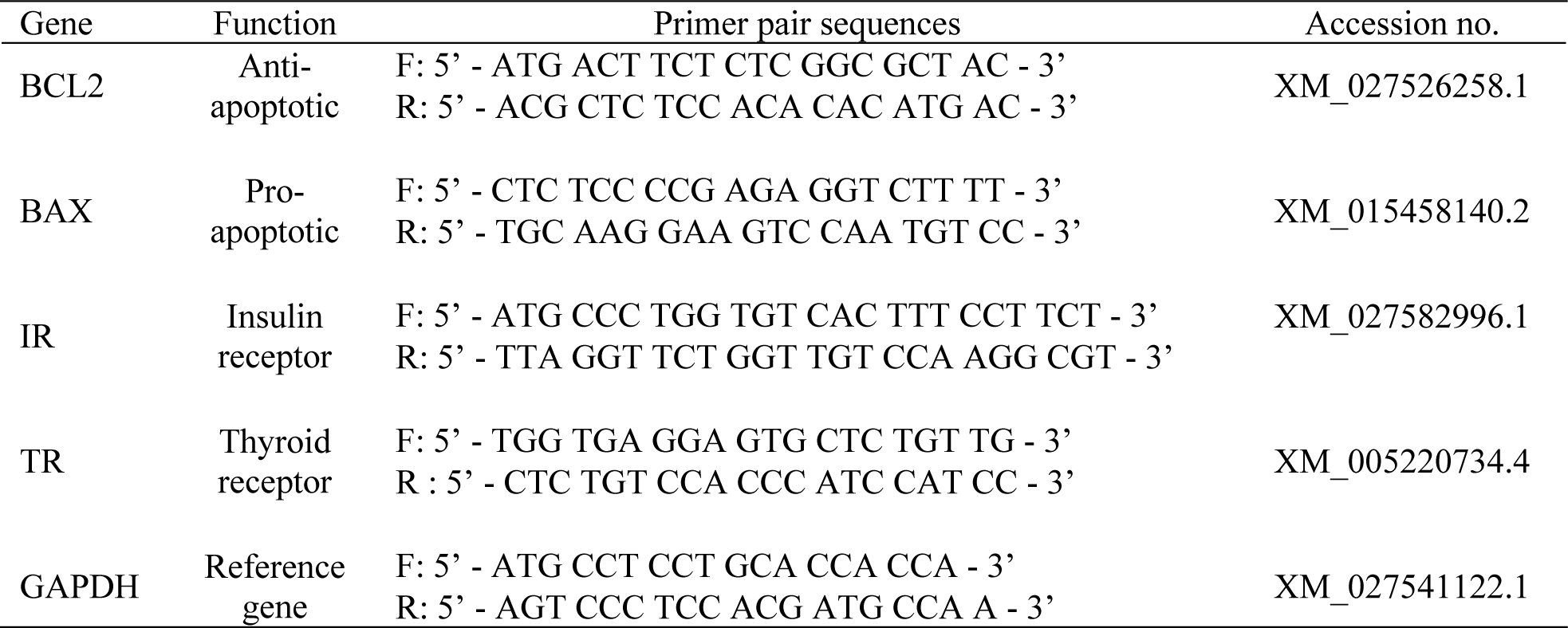
Primer pairs used for Real-Time Reverse-Transcriptase PCR analysis.

### Statistical analyses

Statistical analyses were performed using Sigma Plot version 11 (Systat Software Inc., USA). Normality (Kolmogorov-Smirnov test) and homogeneity of variance (Levene’s tests) were previously evaluated. A comparison of means were analyzed by ANOVA followed by post-hoc test or t-test, when appropriate. Differences were considered significant at P<0.05 (two-sided). Data are presented as percentage or mean (± SEM).

## Results

### Experiment I

#### Follicular survival

Over 2500 preantral follicles were evaluated. The percentage of normal morphological follicles before (non-cultured control) and after 1 or 7 days of culture *in vitro* of preantral follicles is shown (Fig 2). After one day of culture, only ovarian tissues cultured under 10 ng/ml insulin with 1 µg/ml T4 or 10 µg/ml insulin with 0.5 µg/ml T4 showed reduced (P<0.05) normal follicles as compared to non-cultured controls. Follicle survival rates derived from cultures under 10 ng/ml insulin with 1 µg/ml T4 was significantly lower than those cultured with insulin alone. Regardless of insulin and T4 concentrations, the follicle survival was significantly decreased (P<0.05) from day 1 to day 7. On day 7, the combination of 10 ng/ml insulin and 0.5 µg/ml T4 led to higher follicular survival rate than insulin alone. Meantime, only 10 ng/ml insulin and 2 µg/ml T4 combination yielded similar follicular survival rates (P > 0.05) to non-cultured control.

**Fig 2.**
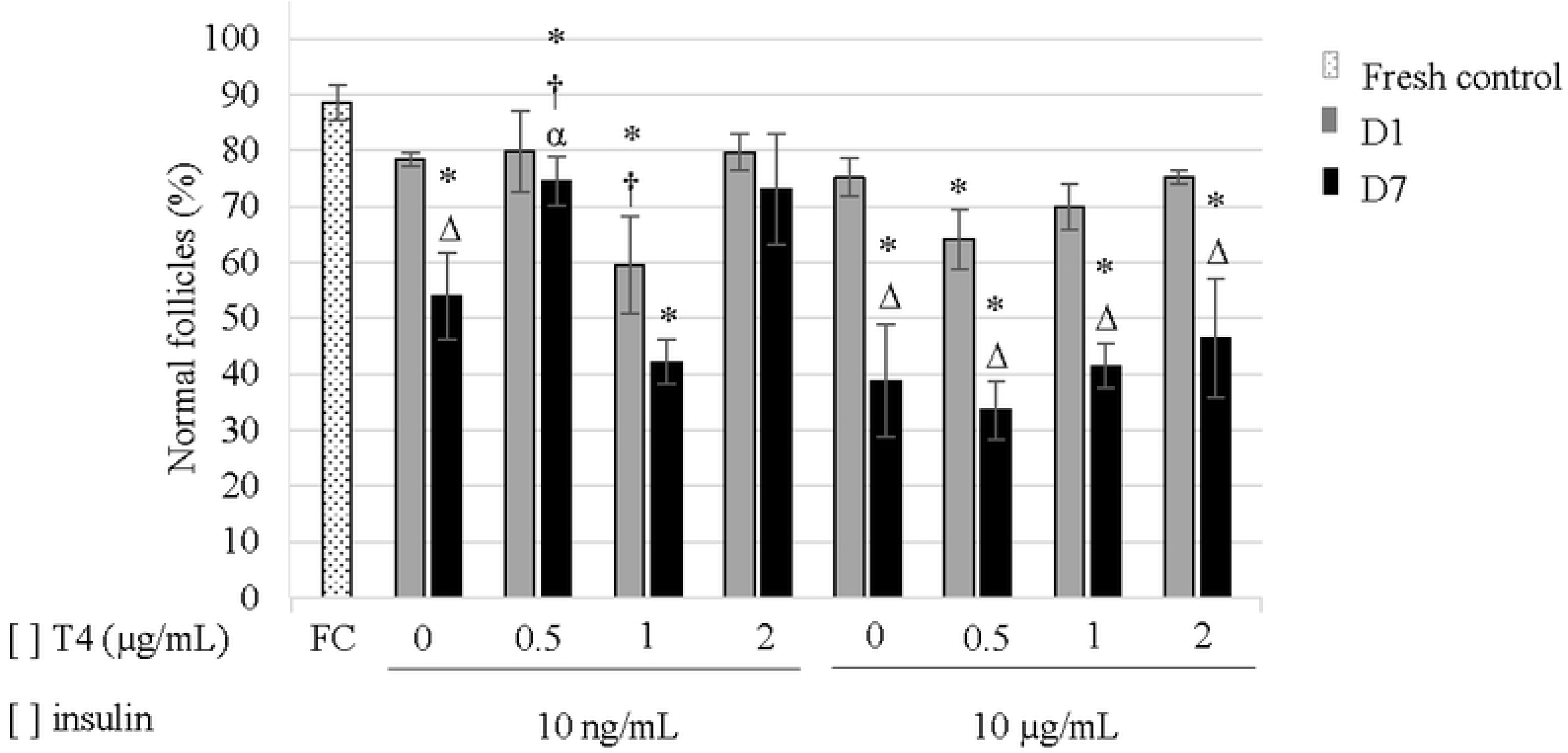
Percentage (mean ± SEM) of morphologically normal preantral follicles. * significant differences compared to non-cultured control. ^†^ significant differences between insulin alone in the same insulin concentration on each day. ^Δ^ significant differences between days in the same treatment. ^α^ significant differences between insulin concentration in the same T4 concentration and on each day. Each symbol indicates a significant difference at P<0.05.

#### Follicular development

The percentage of primordial and developing follicles from non-cultured control and after culture *in vitro* for 1 or 7 days are shown in Fig 3A and Fig 3B, respectively. The most predominant follicle type in non-cultured ovarian tissues was primordial follicle (73.78%) versus 26.22% of developing follicle. In general, ovarian tissues cultured for 7 days led to both, a reduction in the number of primordial follicles and a promotion of follicle development rate (P<0.05) when compared to non-cultured control or ovarian tissues cultured for 1 day. However, on day 7, ovarian tissues cultured under 10 µg/ml insulin with 0.5 or 2 µg/ml T4 had greater follicular development (P<0.05) than their counterparts under 10 ng/ml insulin with the same T4 concentrations.

**Fig 3.**
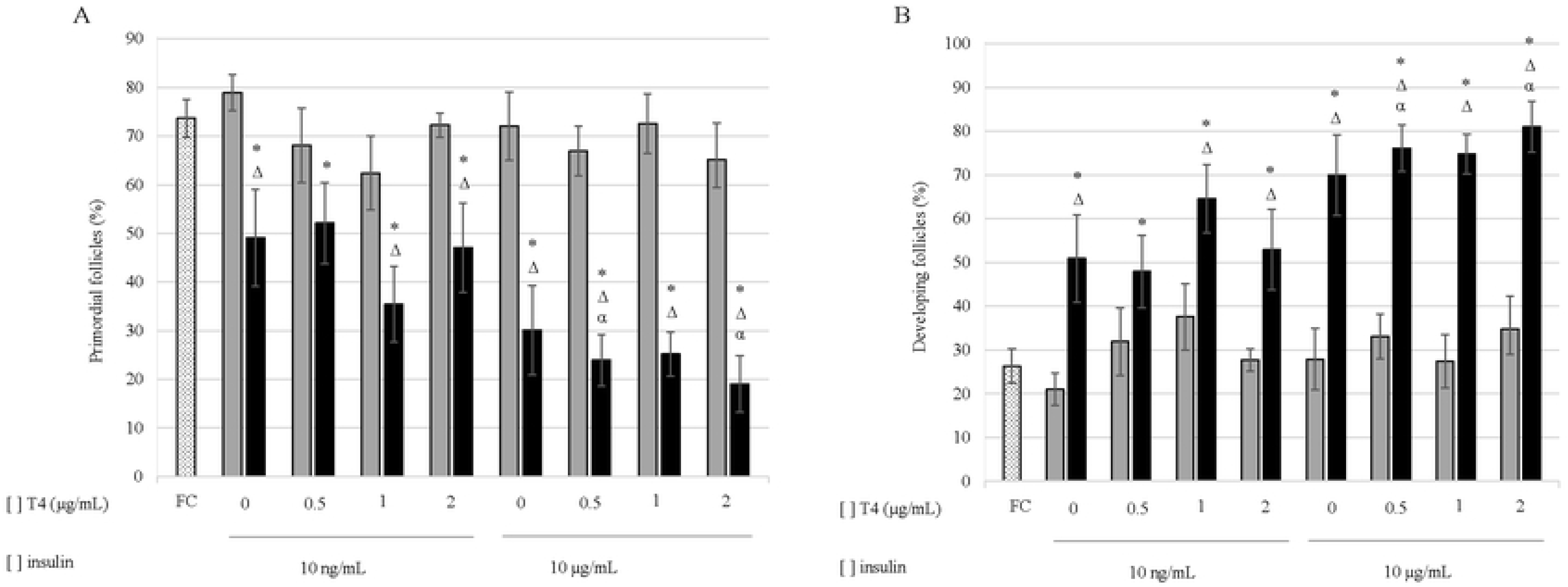
Percentage (mean ± SEM) of primordial (A) and developing follicles (B). * significant differences compared to non-cultured control. ^Δ^ significant differences between days in the same treatment. ^α^ significant differences between insulin concentration in the same T4 concentration on each day. Each symbol indicates a significant difference at P<0.05.

#### Follicle and oocyte diameter

Follicular and oocyte diameters are shown in Table 2. On day 1 of ovarian tissue culture under 10 ng/ml insulin with 2 µg/ml T4, the follicle diameter significantly increased when compared to insulin alone or non-cultured control. Meantime, follicular diameters were significantly increased (P<0.05) under 10 µg/ml insulin with 1 or 2 µg/ml T4, when compared to insulin alone. However, follicles cultured for 1 day under 10 ng/ml insulin with 0.5 µg/ml T4 had better follicle growth (P<0.05) than their counterparts with 10 µg/ml insulin. Under insulin alone (10 ng/ml or 10 µg/ml), follicle diameters were significantly increased (P<0.05) from day 1 to day 7. Only 10 ng/ml insulin led to larger follicle sizes (P<0.05) on day 7, as compared to non-cultured controls. On day 7 of culture, however, culture media supplemented with 2 µg/ml T4 and 10 ng/ml insulin significantly decreased follicle diameters to comparable values with non-cultured controls; while the combination of 2 µg/ml T4 with 10 µg/ml insulin led to significant (P<0.05) larger follicular diameters, when compared to non-cultured controls and any other treatments.

**Table 2.** Goat follicle and oocyte diameters (mean ± SEM).

| | | Follicular diameter ($\mu\text{m}$) | | Oocyte diameter ($\mu\text{m}$) | |
| --- | --- | --- | --- | --- | --- |
| Non-cultured control | | 30.8 ± 0.6 | | 24.5 ± 0.4 | |
| [ ] insulin | [ ] T<sub>4</sub> | Day 1 | Day 7 | Day 1 | Day 7 |
| 10 ng/mL | 0.0 $\mu\text{g/mL}$ | 30.8 ± 0.7 | 33.4 ± 0.9<sup>*Δ</sup> | 24.2 ± 0.5<sup>α</sup> | 25.7 ± 0.6 |
| | 0.5 $\mu\text{g/mL}$ | 32.6 ± 0.9<sup>α</sup> | 31.6 ± 0.6 | 25.9 ± 0.6<sup>*$\dagger\alpha$</sup> | 25.2 ± 0.4 |
| | 1.0 $\mu\text{g/mL}$ | 30.8 ± 0.5 | 32.0 ± 1.1 | 24.1 ± 0.6 | 24.8 ± 0.8 |
| | 2.0 $\mu\text{g/mL}$ | 33.6 ± 0.7<sup>*†</sup> | 29.8 ± 0.6<sup>$\dagger\Delta$</sup> | 25.4 ± 0.7 | 22.8 ± 0.4<sup>*$\dagger\Delta$</sup> |
| 10 $\mu\text{g/mL}$ | 0.0 $\mu\text{g/mL}$ | 30.4 ± 0.6 | 32.5 ± 0.7<sup>Δ</sup> | 22.4 ± 0.6<sup>*</sup> | 25.3 ± 0.6<sup>Δ</sup> |
| | 0.5 $\mu\text{g/mL}$ | 29.9 ± 0.7 | 31.4 ± 0.7 | 22.4 ± 0.5<sup>*</sup> | 24.1 ± 0.6<sup>Δ</sup> |
| | 1.0 $\mu\text{g/mL}$ | 32.3 ± 0.7<sup>†</sup> | 31.3 ± 1.2 | 23.9 ± 0.5<sup>†</sup> | 23.4 ± 1.0<sup>*†</sup> |
| | 2.0 $\mu\text{g/mL}$ | 33.5 ± 1.4<sup>†</sup> | 36.3 ± 0.7<sup>*$\dagger\Delta\alpha$</sup> | 25.5 ± 0.7<sup>†</sup> | 26.2 ± 0.5<sup>*α</sup> |

\* significant differences between fresh control. <sup>†</sup> significant differences between insulin alone, in the same insulin concentration and day. <sup>$\Delta$</sup> significant differences between days in the same treatment. <sup>$\alpha$</sup> significant differences between insulin concentrations, in the same T4 concentration and day. Each symbol indicates a significant difference at P<0.05.

After 1 day of culture, ovarian tissues cultured with 10 ng/ml insulin and 0.5 µg/ml T4 had larger (P<0.05) oocyte than non-cultured controls or under insulin alone. Cultures with 10 µg/ml insulin alone or in combination with 0.5 µg/ml T4 led to significant reduction (P<0.05) of oocyte diameters than their non-cultured control counterparts, and increased oocyte diameter under 10 ng/ml insulin with 0.5 µg/ml T4 (P<0.05). Also on day 1 of culture, ovarian tissues cultured under 10 µg/ml insulin with 1 or 2 µg/ml T4 significantly increased oocyte growth, as compared to insulin alone.

After 7 days of culture, ovarian tissues cultured under 10 ng/ml insulin with 2 µg/ml T4 had lower (P<0.05) oocyte diameters than their counterparts under 10 ng/ml insulin alone, 10 µg/ml insulin with 2 µg/ml T4, or non-cultured control. However, the use of 10 µg/ml insulin alone or in combination with 2 µg/ml T4 led to significant oocyte growth (P<0.05) from day 1 to day 7. The combination of 10 µg/ml insulin with 1 µg/ml T4 significantly reduced oocyte diameter on day 7, when compared to insulin alone or non-cultured controls. Only ovarian tissues cultured under 10 µg/ml insulin with 2 µg/ml T4 had greater (P<0.05) oocyte growth than non-cultured controls.

#### Stromal cell density

Stromal cell densities (number of cells per 100 µm^2^) of non-cultured control (Fresh control) and cultured (1 or 7 days) ovarian tissues are shown in Fig 4. After 1 day of culture and regardless of the insulin concentrations, culture supplementations with 0.5 µg/ml T4 significantly (P<0.05) increased the number of stromal cells as compared to both non-cultured control and insulin alone, and culture supplementations with 2 µg/ml T4 reached similar increase. The presence of 2 µg/ml T4 yielded greater (P<0.05) stromal cell numbers when associated to 10 ng/ml rather than 10 µg/ml insulin. Surprisingly, after 1 day of culture, ovarian tissues cultured under 10 ng/ml insulin with 1 µg/ml T4 led to significant reduction (P<0.05) of the stromal cell density.

**Fig 4.**
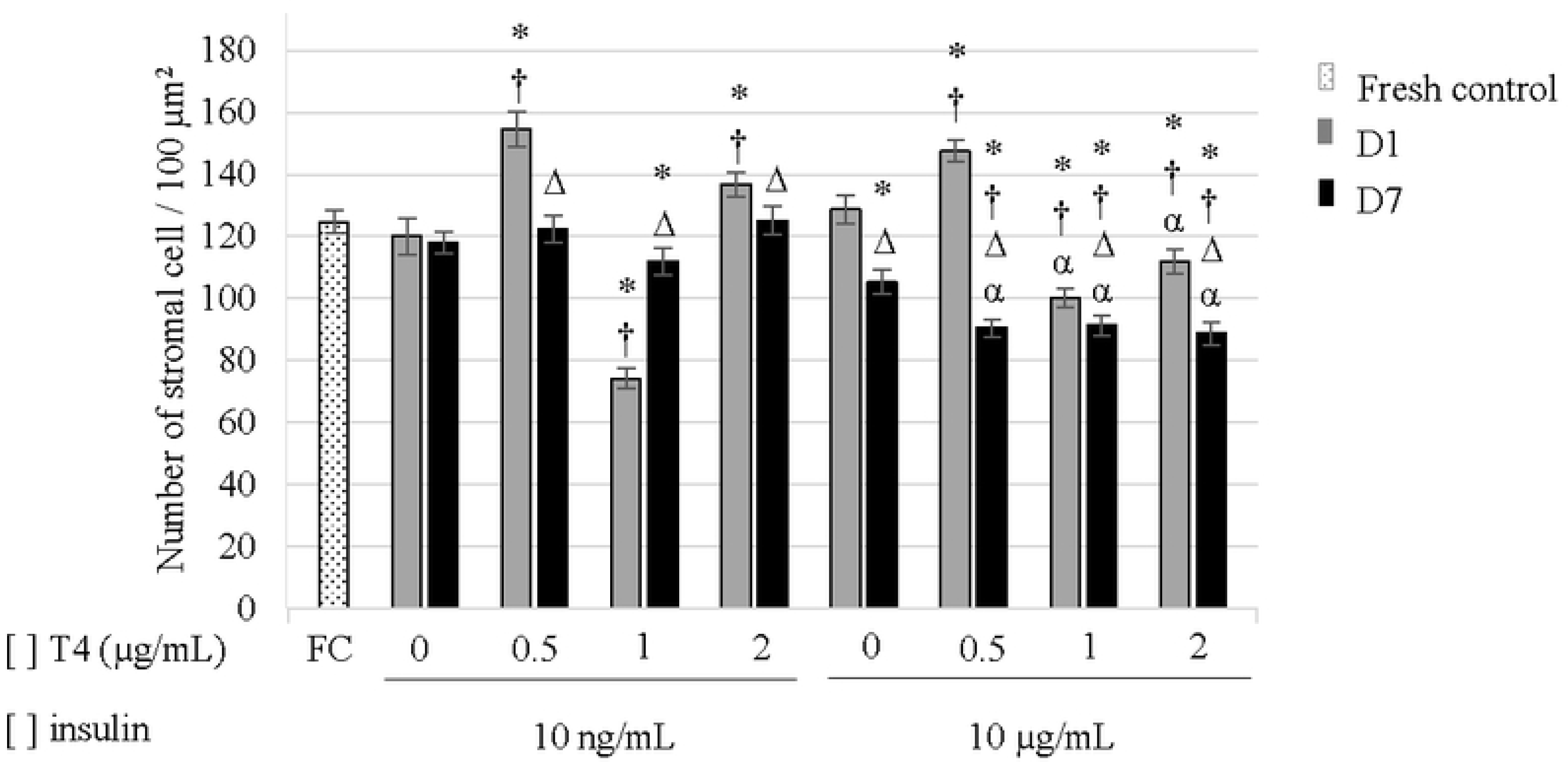
Mean (± SEM) density of caprine ovarian stromal. * significant differences compared to non-cultured control. ^†^ significant differences between insulin alone in the same insulin concentration on each day. ^Δ^ significant differences between days in the same treatment. ^α^ significant differences between insulin concentration in the same T4 concentration on each day. Each symbol indicates a significant difference at P<0.05.

Under 10 µg/ml insulin, there was a global and significant decrease of stromal cell density after 7 days of culture, as compared to day 1 or non-cultured controls. In contrast, cultures under 10 ng/ml insulin had better stromal cell density than 10 µg/ml insulin, regardless of T4 concentrations. The combination of 10 ng/ml insulin with 0.5 or 2 µg/ml T4 significantly reduced (P<0.05) the number of stromal cells from day 1 to day 7, while these numbers were significantly increased (P<0.05) with 1 µg/ml T4, but remained lower (P<0.05) than their non-cultured control counterparts.

#### Estradiol production

The Fig 5 shows estradiol production by ovarian cortex cultured in situ for one day. Due to the large individual variation among ovarian fragments, differences in the levels of estradiol production were not observed among the treatment groups. However, when data were taken together into two groups (10 ng/ml vs. 10 µg/ml insulin, regardless of T4 concentrations), ovarian fragments cultured with 10 ng/ml insulin showed higher estradiol production (P<0.05) than fragments cultured in 10 µg/ml insulin. On day 7, the estradiol levels in all treatment groups were under the detection limit of the assay (9 pg/ml).

**Fig 5.**
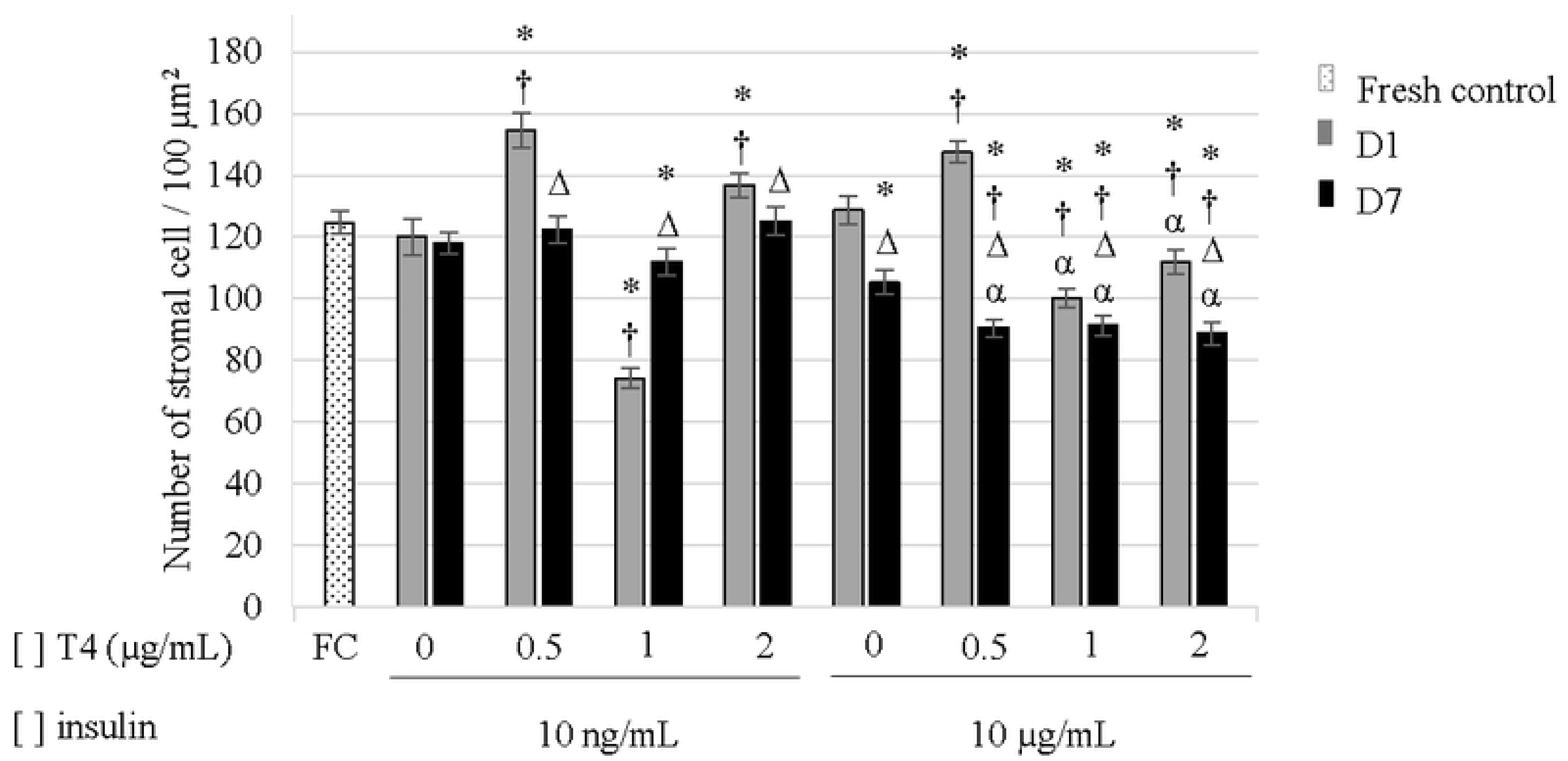
Estradiol production (mean ± SEM) by ovarian tissue cultured for 1 day. * significant differences between treatments (P<0.05).

#### ROS production

The ROS production in ovarian tissues cultured *in vitro* for 1 or 7 days is shown (Fig 6). Unlike treatment groups containing 10 ng/ml insulin, all treatments with 10 µg/ml insulin significantly decreased the ROS levels from day 1 to day 7. On the other hand, ovarian tissues cultured under 10 ng/ml insulin with 0.5 or 2 µg/ml T4 had increased (P<0.05) ROS production from day 1 to day 7. Finally, cultures under 10 ng/ml insulin with 0.5 or 1 µg/ml T4 displayed higher levels of ROS (P<0.05) than their counterparts under 10 µg/ml insulin with any T4 concentrations.

**Fig 6.**
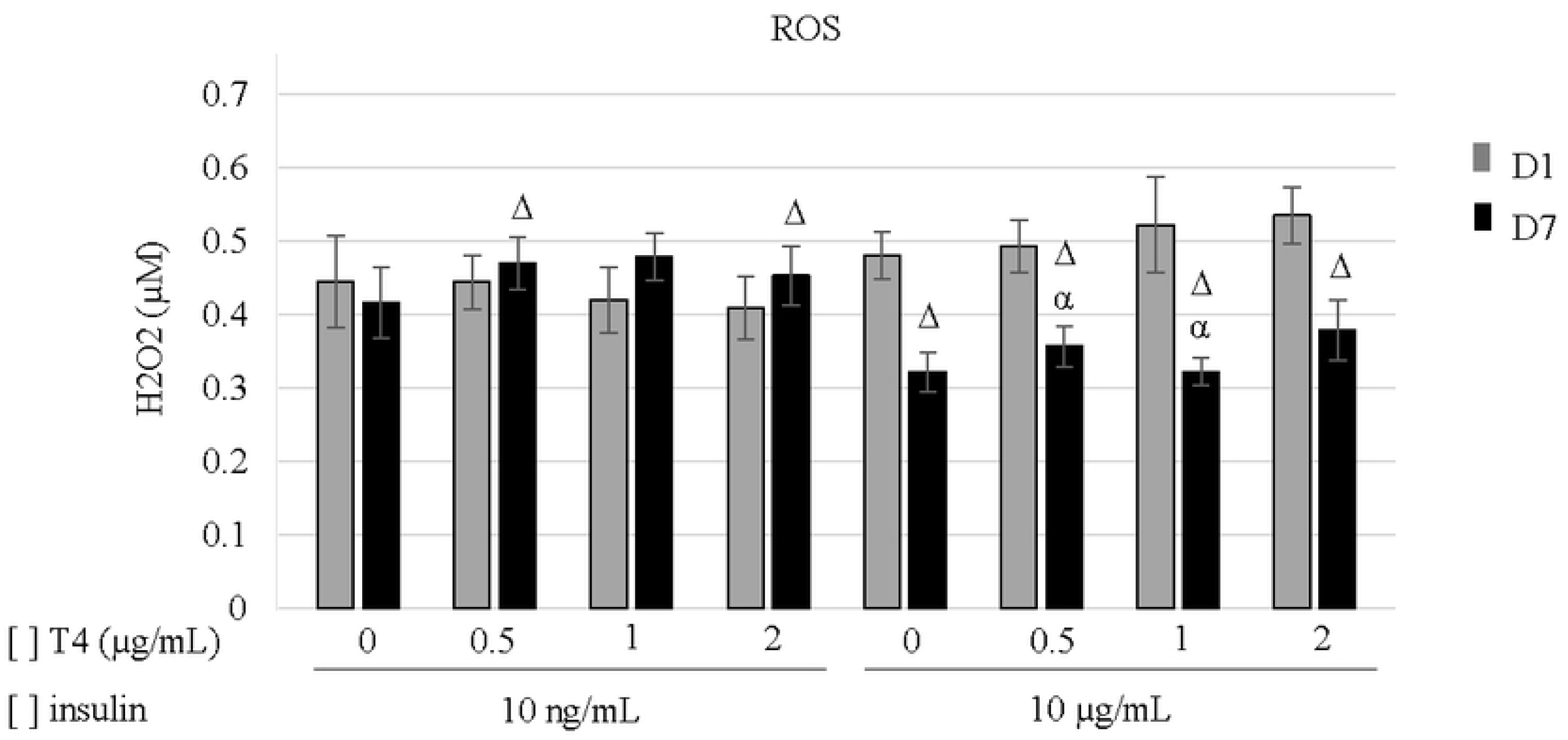
ROS production (mean ± SEM) of ovarian tissue cultured for 1 or 7 days. ^Δ^ significant differences between days in the same treatment (P<0.05). ^α^ significant differences between insulin concentration in the same T4 concentration on each day (P<0.05).

### Experiment II

#### Relative gene expression

Transcript levels related to both thyroid and insulin hormone receptor (TR/IR), and apoptosis (*Bcl2/Bax*) evaluated on day 7 of culture are shown in Fig 7. Overall, ovarian tissues cultured under 10 µg/ml insulin, regardless of T4 concentrations led to reduced (P<0.05) TR expression levels, when compared to non-cultured control (Fig 7A). Under 10 ng/ml insulin, the TR expression did not differ (P > 0.05) from non-cultured control (FC) versus insulin alone or combined with 2 µg/ml T4 (Fig 7A). Compared to non-cultured controls, the presence of 10 µg/ml insulin significantly decreased IR expression, regardless of T4 concentrations (Fig 7B). Under 10 ng/ml insulin, there was a significant reduction of IR expression by the presence of T4 (P<0.05), as compared to insulin alone or non-cultured controls (Fig 7B). In comparison to non-cultured controls, the *Bcl2/Bax* ratios were decreased in all treatments during cultures (Fig 7C), but cultures under 10 µg/ml insulin exhibited lower ratios than 10 ng/ml insulin, regardless T4 concentrations.

**Fig 7.**
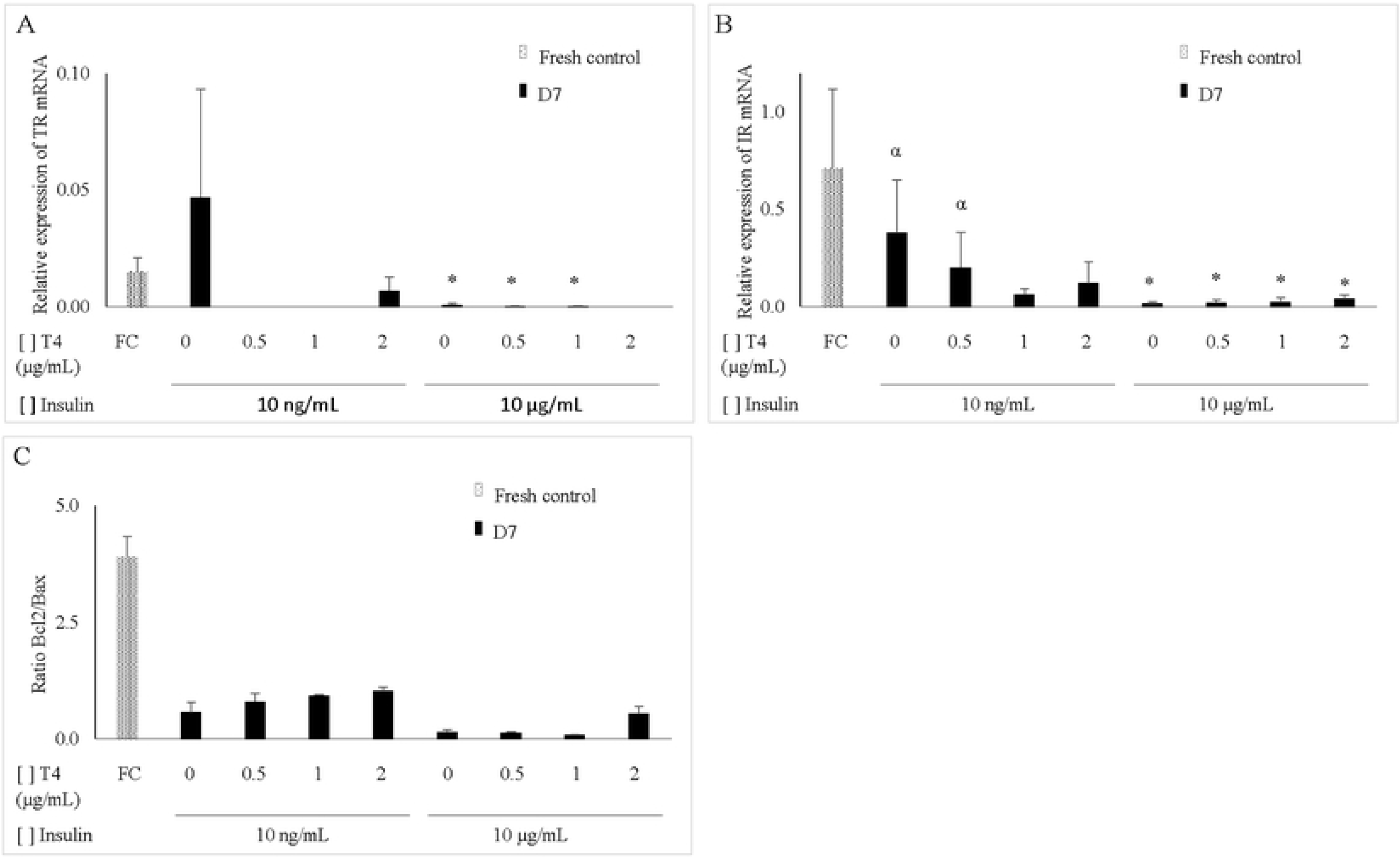
Relative mean (± SEM) expression of *TR* (A), *IR* (B), and ratio *Bcl2/Bax* (C). * significant differences between fresh control (P<0.05). ^α^ significant differences between insulin concentration in the same T4 concentration (P<0.05).

## Discussion

To our knowledge, this is the first study to investigate the role of T4 in different concentrations (0.5, 1 or 2 µg) during *in vitro* culture of primordial follicles in the presence of low (10 ng/ml) or high (10 µg/ml) insulin levels. Herein, we observed concentration-dependent effects of both hormones on follicle survival and development.

Thyroid hormones are essential for female reproductive function, and a dysregulation of these hormones have been associated with infertility (3). Indeed, the subtle balance of T4 levels during *in vitro* culture of primordial follicles affected the follicular dynamic. Herein, we observed that, unlike the treatments with 10 µg/ml insulin, the addition of T4 to a basic medium containing 10 ng/ml insulin maintained follicle survival from day 1 to day 7. Moreover, it was demonstrated that the combination of 0.5 µg/ml T4 and 10 ng/ml insulin enhanced follicle survival in comparison to T4-free treatment (i.e., 10 ng/ml insulin alone) or when associated to high levels of insulin. It is known that insulin increases glucose uptake by translocation of GLUT (32) and THs could improve this effect (33), which reduces the amount of glucose in the culture medium (34). However, our culture system uses a fixed glucose concentration (1 g/l), and thus, low levels of combined insulin and thyroxine could be more appropriate to maintain the nutritional support to the survival of ovarian tissue.

In the present study, treatments (i.e., 10 ng/ml insulin + T4 at 0.5 or 2 µg/ml) that provide the best outcomes for follicular survival exhibited higher ROS production from day 1 to day 7 of culture. ROS are products of normal metabolism and can be beneficial to cells acting as redox messenger in intracellular signaling (35). Recently, our team demonstrated an association between ROS levels and follicle survival (36). Although the cultured tissues exhibited metabolic activity represented by ROS production, our results showed a reduction of *Bcl2/Bax* mRNA ratio at the end of culture, regardless of treatment. However, *Bcl2/Bax* ratio was lower in the treatments using high insulin than treatments using low insulin. Bcl2 and Bax proteins act as a pro-cell survival or pro-cell death, respectively in the apoptosis pathway (37). Thus, a reduction in the *Bcl2/Bax* rate indicates the activation of apoptosis signaling. Here we used caprine ovarian cortex tissues that are filed with more stromal cells than ovarian follicles (38), and the decrease in *Bcl2/Bax* ratio could be due to the increase in stromal cell apoptosis.

In general, media containing low insulin were better than those with high insulin for the maintenance of stromal cell density, which functionality remains for demonstration. Indeed, estrogen that is also produced in stroma cells (39) was not detected in the culture medium, suggesting a possible stromal cell apoptosis occurrence. On the other hand, after one day of culture, low insulin treatments led to more estrogen production than high insulin, which corroborates with follicle survival and highlights the importance of insulin to glucose balance on estrogen producing ovarian cells (granulosa and theca cells).

All treatments improved the percentage of developing follicles, and better results were obtained with high levels thyroxine and insulin that yielded greatest follicle and oocyte diameters. The association of thyroid hormones with growth factors and/or hormones enhance follicle growth (12, 17) and granulosa cell proliferation (33, 40). However, the requirement of high levels of thyroxine for follicular development may be due to the presence of BSA in the culture medium. Indeed, thyroid hormones can bind seric/plasmic albumin (41), constituting an *in vitro* obstacle for the maintenance of free thyroid hormone fractions in the culture medium. Thyroid receptors were detected for the first time in caprine ovarian tissues. Decreased expression of thyroid and insulin receptors were observed under high level insulin conditions, which could be explained by a possible increase of their translation into proteins, enhancing glycolytic activity that delivers necessary energy for follicle development (33, 42).

In conclusion, both survival and development of primordial follicles in culture were differently affected by insulin and thyroxine concentrations. The combination of insulin and thyroxine at lower concentrations his beneficial for follicular survival. High levels of combined insulin and thyroxine had better outcomes in follicle development.

## Author contributions

### Conceptualization

Victor M. Paes, Jose R. Figueiredo, Jean M. Feugang.

### Formal analysis

Benner G. Alves.

### Investigation

Victor M. Paes, Laritza F. Lima, Anna-Clara A. Ferreira, Carlos H. Lobo, Benner G. Alves, Ana-Paula R. Rodrigues, Ariclecio C. Oliveira.

### Methodology

Victor M. Paes, Ariclecio O. Oliveira, Jose R. Figueiredo, Jean M. Feugang.

### Resources

Ariclecio O. Oliveira, Ana-Paula R. Rodrigues, Jose R. Figueiredo, Jean M. Feugang.

### Supervision

Jose R. Figueiredo, Jean M. Feugang.

### Validation

Victor M. Paes.

### Writing – original draft

Victor M. Paes, Jose R. Figueiredo, Jean M. Feugang.

### Writing – review & editing

Victor M. Paes, Jose R. Figueiredo, Jean M. Feugang.

